# Obesity modifies cell fate plasticity of *Pdgfrα-*expressing adipocyte progenitors to promote an aberrant mammary microenvironment

**DOI:** 10.1101/2023.12.28.573587

**Authors:** Sharon A. Kwende, Prashant Nuthalapati, Jacqulene P. Sunder Singh, Subhajit Maity, Dun Ning, Douglas G. Shryock, Zhenyu Xuan, Purna A. Joshi

## Abstract

Excess fat gain culminating in obesity is a mounting global pandemic associated with a myriad of diseases including a higher risk of developing multiple cancers such as breast cancer. The underlying biological basis for obesity-linked breast cancer is attributed to alterations in adipose tissue-derived endocrine, metabolic and inflammatory factors. However, the precise cell types responsible for generating a cancer-susceptible mammary gland in obesity are not well understood. Using diet-induced obesity in conjunction with genetic reporter mouse models, we reveal elevated mammary adipocyte progenitors (MAPs) expressing Platelet-Derived Growth Factor Receptor alpha (PDGFRα) in the mammary tissue microenvironment during obesity. Single-cell RNA sequencing analysis demonstrate intensified obesity-driven trajectories of MAPs to immune and epithelial progenitor cells. Further, lineage tracing indicates an invasive cellular state of MAPs that confers greater access of MAP descendants into the mammary epithelium concomitant with their transition into immune and epithelial cell fates. MAPs in obesity exhibit heightened activation of cancer-associated and inflammatory pathways where *Egfr* is identified as a common target upregulated in MAPs. Mechanistic studies on purified MAPs with a major obesity diet component and EGFR stimulation corroborate dynamic EGFR-mediated MAP responses. Our findings uncover MAPs as key contributors to forging an aberrant mammary gland in obesity, providing insight into the potential utility of targeting this cell lineage for eradicating cancer risk related to augmented adiposity.

## Introduction

Beyond health issues related to metabolic syndrome, excess body fat increases the risk of at least 13 types of cancer including breast cancer^1^. Obesity is generally linked to increased risk of hormone receptor-positive breast cancer in post-menopausal women^2^, although studies also report a positive association with premenopausal breast cancer risk especially in relation to the aggressive hormone receptor-negative, HER2^-^ (triple-negative) subtype^3–6^. The prevalence of obesity is also greatest in women of African ancestry who are disproportionately at higher risk of aggressive triple-negative breast cancer compared to other racial groups^7,8^. Mechanisms involving estrogen, adipokines, insulin, growth factors, and inflammation are proposed to underlie the pathophysiological link between increased adiposity and breast cancer^1,9^.

Adipose tissue expansion results from hypertrophy and hyperplasia of adipocytes, where the latter is facilitated by adipocyte progenitors^10^. Adipocyte progenitors marked by PDGFRα that can give rise to functional adipocytes *in vivo* have been described^11–13^, but their activity may become dysregulated in obesity^14,15^. We have previously reported PDGFRα^+^ mammary adipocyte progenitors (MAPs) in the stromal microenvironment of mammary tissue that transition into epithelial lineages during expansion of the murine mammary epithelium. The contribution of MAPs to shaping a mammary gland conducive to tumorigenesis in obesity is not known. In this study, we sought to fill this critical gap by leveraging a diet-induced obesity mouse model together with MAP reporter, lineage tracing strains and single-cell RNA sequencing. In both wild-type and genetic *Pdgfrα* MAP reporter mice fed a high-fat diet, we find a remarkable increase in mammary tissue MAPs. Single-cell RNA sequencing (scRNA-seq) of GFP^+^ MAP reporter cells identify enhanced cell fate trajectories of a MAP subset to immune, basal and luminal epithelial progenitor populations. In PDGFRα^+^ cell tracing studies, MAP descendants in obesity are observed to express epithelial and immune markers, and penetrate the mammary epithelial space in the absence of an intact basement membrane. Delving into obesity-mediated transcriptomic changes in MAPs expose an enrichment of cancer-associated and inflammatory pathways following a high-fat diet, where *Egfr* is found to be highly expressed. EGFR stimulation of isolated GFP^+^ MAPs from reporter mice along with palmitic acid, a major high-fat diet component, leads to pronounced proliferation and differentiation of MAPs. Our results underscore the presence of a divergent EGFR-activated MAP cell population that engenders a cancer-favorable mammary milieu in obesity.

## Results

### PDGFRα^+^ MAPs expand in the mouse mammary stroma upon high fat diet-induced obesity

We first investigated MAP distribution in the mammary gland during obesity. Adult 8- to 12-week-old C57BL/6J female mice were fed a high-fat diet (HFD) with 60% kcal from fat^16^ or a matched control diet (CD) for 8-10 weeks **(Fig. 1a)**. HFD mice were observed to gain 25% more weight (p<0.01) by 8 weeks compared to CD mice, and exhibited significantly elevated glucose and insulin levels **(Fig. 1b-d)** indicative of obesity. Whole mounts of mammary glands isolated from HFD-induced obese mice showed an enlarged fat pad and a greater branching network compared to CD mice accompanied by increased size, number of branch points and mammary gland mass **(Fig. 1e-h)**. Immunostaining analysis of mammary tissues showed a characteristic stromal-specific expression pattern for PDGFRα^+^ MAPs with absence of its expression in the EpCAM^+^ mammary epithelium as previously observed^17^ **(Fig. 1i)**. Of note, PDGFRα^+^ MAPs had a more prominent presence in glands from HFD mice compared to controls. To quantify MAPs, flow cytometry was performed using previously published cell surface markers^17^ to discern the luminal epithelial (Lin^-^ CD49f^lo^ EpCAM^+^), basal epithelial (Lin^-^ CD49f^hi^ EpCAM^+^), stromal (Lin^-^ CD49f^-^ EpCAM^-^) and lineage^+^ immune, endothelial and erythrocyte (Lin^+^ CD45^+^ CD31^+^ Ter119^+^) mammary subpopulations **(Fig. 1j)**. Upon analysis with a fluorescence dye-conjugated antibody to the cell surface receptor, PDGFRα^+^ cells were not present in luminal, basal and Lin^+^ subsets in HFD or CD glands, and were exclusively detected in the stromal subset **(Fig. 1j and Supplementary Fig. 1)**. Importantly, the proportion of PDGFRα^+^ MAPs increased by 33% in obese mice compared to controls **(Fig. 1j, k)**. These results inferred an activated MAP population in the mammary microenvironment during obesity.

**Figure 1.**
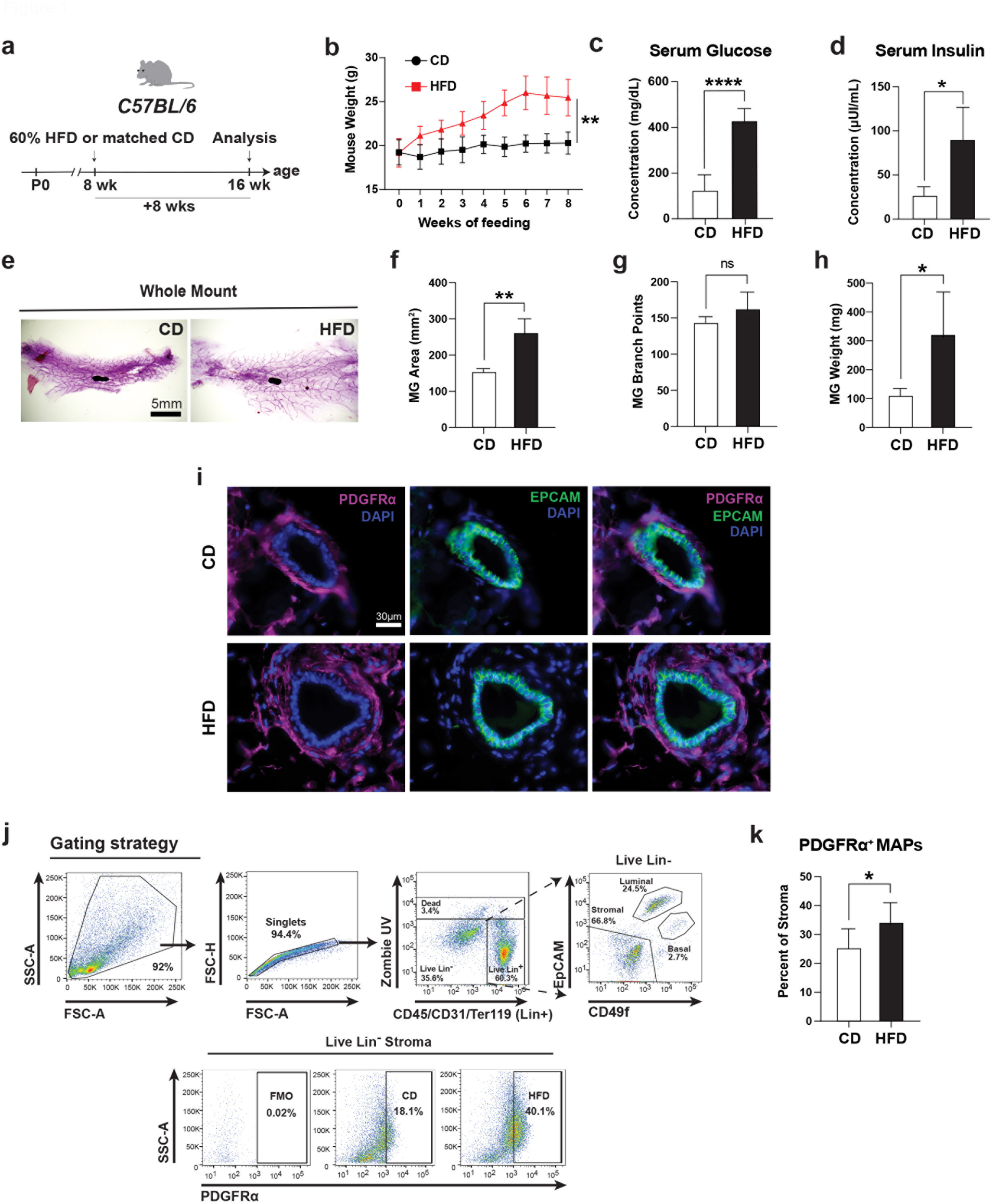
Mammary adipocyte progenitors (MAPs) marked by PDGFRα are elevated in the mouse mammary gland upon diet-induced obesity. **(a)** High-fat diet (HFD) and matched control diet (CD) regimen in wild-type C57BL/6J female mice. **(b)** Mouse weight gain over an 8-week period of HFD feeding compared to control (n=7 per group). **(c)** Serum glucose and **(d)** Insulin levels in HFD compared to CD (n=6 per group). **(e)** Representative whole mounts of HFD and CD wild-type mammary glands. **(f)** Mammary gland area, **(g)** Branch points per mammary gland, and **(h)** Mammary gland mass of HFD and CD mice (n=4 per group). **(i)** Immunofluorescence for PDGFRα and epithelial marker EpCAM on representative HFD and CD mammary tissue sections (n=7 mice per group, 4 fields per section). **(j)** Flow cytometry gating strategy and analysis of PDGFRα^+^ MAPs in mammary stroma of HFD and CD mice; Fluorescence Minus One (FMO) control for gating PDGFRα^+^ cells. **(k)** Quantification of PDGFRα^+^ MAPs in stroma shown in (j) (n=7 mice per group). Data represent mean ± s.d. *P< 0.05, **P<0.01, ****P<0.0001.

### A genetic *Pdgfra* reporter affirms abnormal MAP activity in obesity

Next, we utilized the *Pdgfrα* genetic reporter strain, *Pdgfrα^H2B-eGFP^*, which harbor a histone H2B-eGFP fusion reporter controlled by the endogenous *Pdgfrα* promoter, enabling us to observe and enumerate MAPs as H2B-eGFP^+^ cells in the mammary gland. Adult *Pdgfrα^H2B-eGFP^* females were fed a 60% kcal HFD to induce obesity or a CD diet **(Fig. 2a)**. Similar to wild-type mice, *Pdgfrα^H2B-eGFP^*mice subjected to a HFD showed significant weight gain, blood insulin and glucose levels, and larger mammary glands **(Fig. 2b-f)**. In immunostained mammary tissue sections from obese mice, cell proliferation detected with the Ki67 marker was greater in GFP^+^ MAPs compared to controls **(Fig. 2g)**. When mammary cells were analyzed by flow cytometry, GFP^hi^ cells, and GFP^lo^ cells with a 19-fold lower mean fluorescence intensity on average than GFP^hi^ cells were observed in mammary subpopulations **(Fig. 2h-k)**. In particular, GFP^lo^ cells were seen in luminal, basal and Lin^+^ subsets, whereas GFP^hi^ cells representative of MAPs were present in the CD49f^-^ EpCAM^-^ stromal compartment. Since PDGFRα^+^ cells are stromally-localized, GFP^lo^ cells are likely progeny of GFP^hi^ MAPs akin to that observed for Lgr5-GFP^lo^ progenitors from Lgr5-GFP^hi^ stem cells in the intestine^18^. As observed with flow cytometry analysis for cell surface PDGFRα in obese wild-type mice, GFP^hi^ MAPs from *Pdgfrα^H2B-eGFP^* glands were strikingly elevated 1.6-fold in the mammary stroma of obese versus lean mice **(Fig. 2i, l)**. Interestingly, GFP^lo^ Lin^+^ cells significantly increased in HFD, while the proportion of luminal and basal cells were reduced in HFD, likely due to the relative expansion in the stromal compartment **(Fig. 2m-o)**. Thus, data from the genetic *Pdgfrα* reporter model confirms the obesity-triggered increased proliferative activity of MAPs.

**Figure 2.**
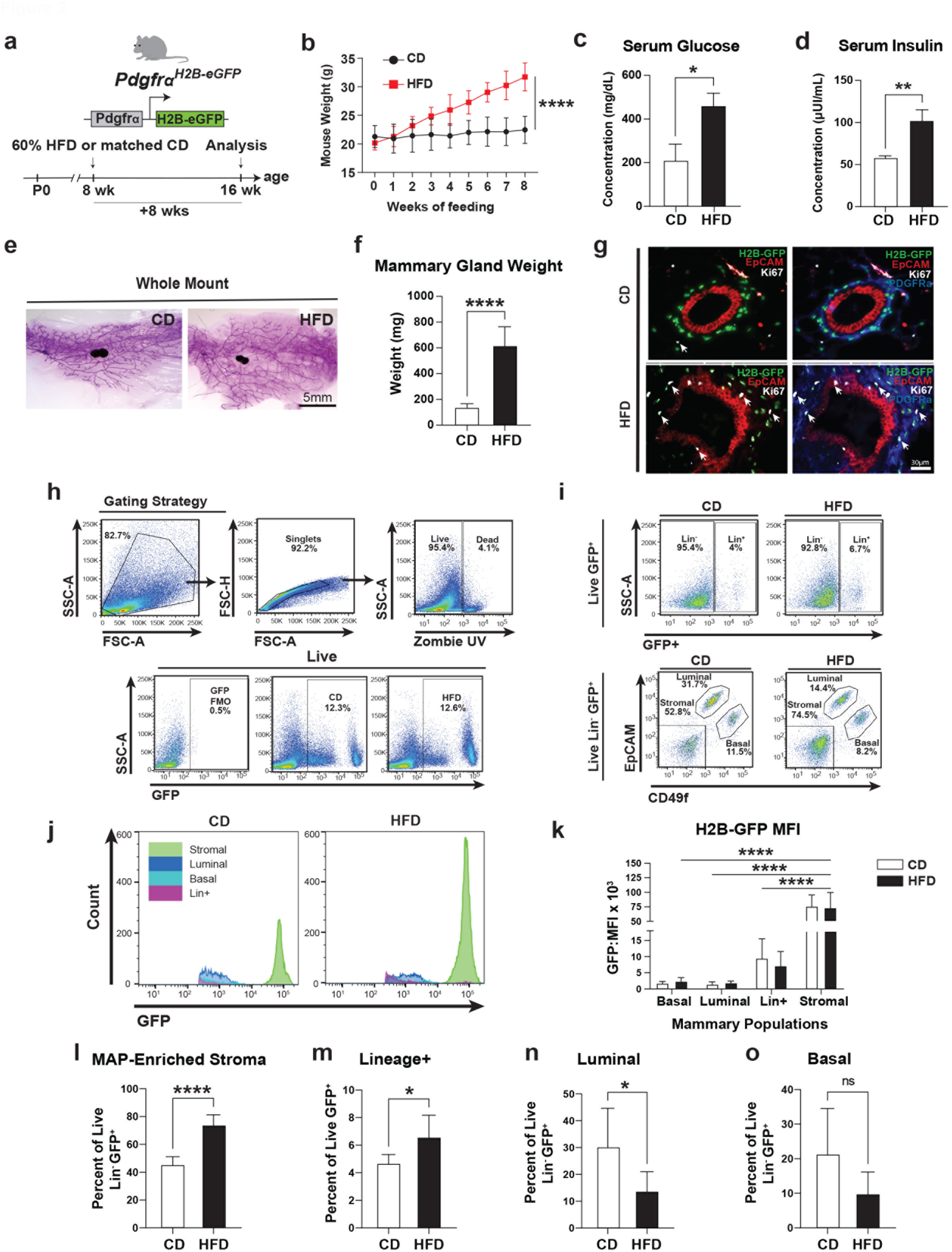
*Pdgfra* H2B-eGFP-expressing MAPs in genetic reporter mice exhibit increased proliferation concomitant with their expansion in obesity. **(a)** *Pdgfrα^H2B-eGFP^* reporter mice were fed an obesity-inducing HFD or matched CD. **(b)** Mouse weight over a period of 8 weeks (n=7 for HFD and n=5 for CD). **(c)** Serum glucose and **(d)** Serum insulin for CD versus HFD MAP reporter mice (n=3 per group). **(e)** Whole mounts representative of mammary glands isolated from reporter mice fed HFD or CD. **(f)** Weight of mammary glands from CD and HFD mice (n= 7 for HFD and n=5 for CD). **(g)** Immunofluorescence detection of Ki67^+^ cell proliferation in *Pdgfrα^H2B-eGFP^* mammary tissue sections co-stained for H2B-GFP reporter, PDGFRα and EpCAM; white arrows indicate proliferating cells. **(h)** Gating strategy for flow cytometry analysis of H2B-GFP^+^ cells in dissociated mammary tissues from HFD and CD mice. **(i)** Representative flow cytometry plots of GFP^+^ cells in Lineage^+^ (CD45^+^ CD31^+^ Ter119^+^; Lin^+^) population and Lineage^-^ (Lin^-^) luminal, basal and MAP-enriched stromal cells from CD and HFD mammary glands. **(j)** Representative histograms depicting GFP mean fluorescence intensity (MFI) on the x-axis and cell counts on the y-axis for distinct cellular subsets. **(k)** MFI values for GFP in the different mammary subpopulations (n=7 for HFD and n=5 for CD). Quantification of H2B-GFP^+^ cells in **(l)** Lin^-^ MAP-enriched stromal subset, **(m)** Lin^+^ subpopulation that includes immune and endothelial cells, **(n)** Lin^-^ luminal subpopulation and **(o)** Lin^-^ basal subpopulation. For all graphs, error bars represent s.d. *P< 0.05, **P<0.01, ****P<0.0001.

### Obesity drives altered cell fate trajectories of MAPs

To determine the transcriptomic profile and cellular state of MAPs in obesity, we performed scRNA-seq on fluorescence-activated cell sorting (FACS)-isolated GFP^+^ mammary cells which included GFP^hi^ and GFP^lo^ cells from both HFD- and CD-fed *Pdgfrα^H2B-eGFP^* mice **(Fig. 3a)**. Following data preprocessing and quality control **(Supplementary Fig. 2)**, analysis of 15,698 GFP^+^ cells identified a total of 11 clusters representing distinct cell types including MAPs, and interestingly, basal cells, luminal progenitors, immune cells, and endothelial cells based on our curated gene marker set **(Fig. 3b, c and Supplementary Fig. 3a)**. MAPs comprised of 6 clusters which specifically expressed *Pdgfrα*, in addition to *Cd34* and *Ly6a* which are established adipocyte progenitor markers^11,19^. Two clusters of luminal progenitors (LP1, LP2) expressing *Krt18* and luminal progenitor markers *Kit*, *Elf5* and *Aldh1a3* were ascertained with LP2 having higher expression of *Elf5* while LP1 showed greater *Aldh1a3* expression. Basal, endothelial and immune clusters had specific expression of *Krt14*, *Pecam1* and *Ptprc* respectively. We then employed the CytoTRACE^20^ computational tool to discern the hierarchical differentiation potential between different clusters in our scRNA-seq data **(Fig. 3d)**. Notably, MAPs exhibited a higher CytoTRACE score globally, suggesting a more stem-like phenotype, followed by LP2, LP1, basal, immune and endothelial cells which had progressively lower predicted CytoTRACE scores reflecting cell populations with increasing differentiation.

**Figure 3.**
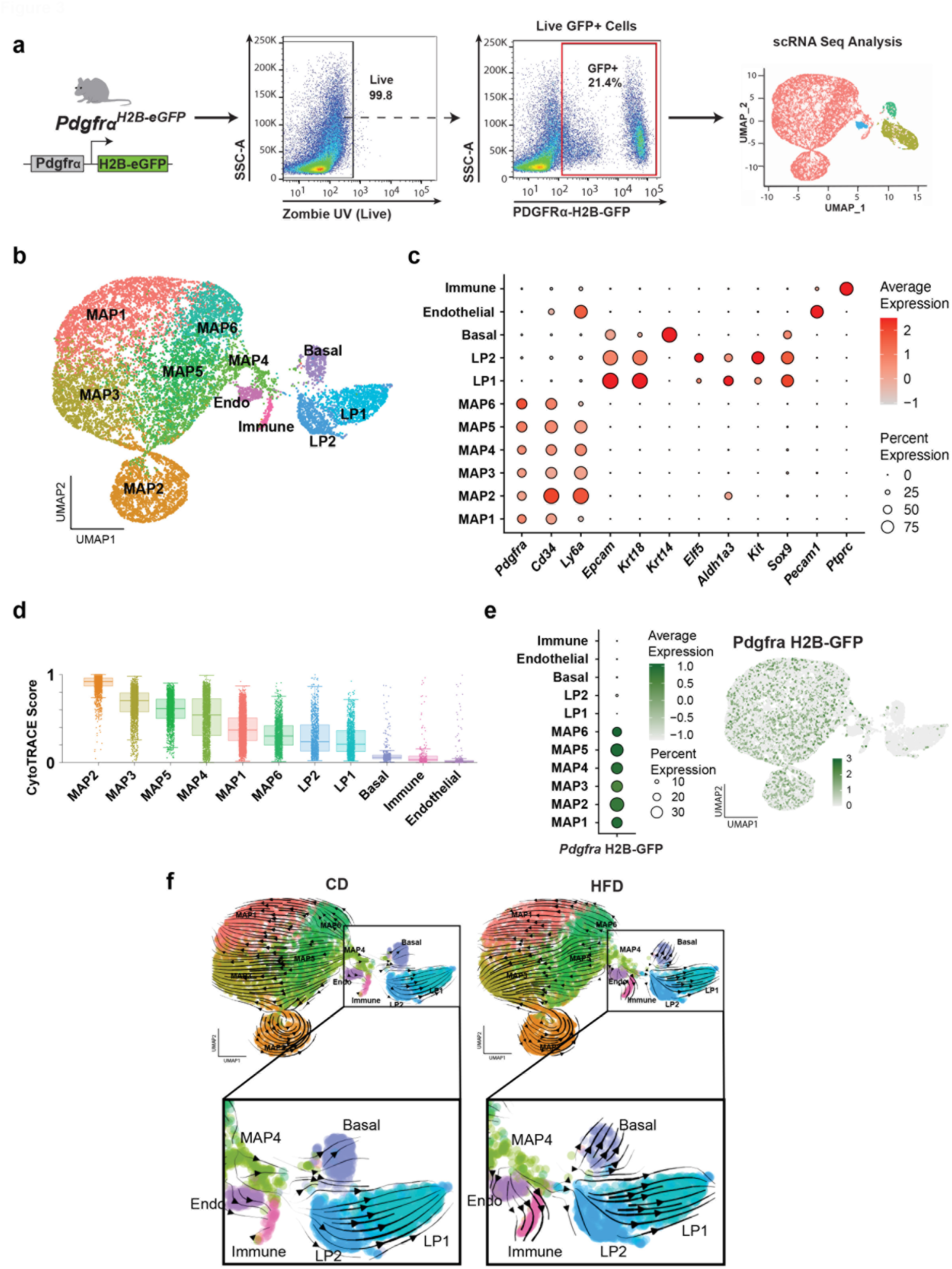
Single cell analysis of the *Pdgfra* H2B-eGFP^+^ lineage expose intensified cell fate trajectories of MAPs to epithelial and immune cells in obesogenic conditions. **(a)** Live H2B-eGFP^+^ cells from *Pdgfrα^H2B-eGFP^* reporter mammary glands of HFD and CD mice were FACS-sorted and prepared for scRNA-seq. **(b)** Uniform manifold approximation and projection (UMAP) visualization of H2B-eGFP^+^ mammary cells combined from HFD and CD, representing 15,698 annotated scRNA-seq profiles, and 11 total clusters: 6 MAP, 2 luminal progenitor (LP) epithelial, 1 basal epithelial, 1 endothelial, and 1 immune. **(c)** Dot plot of representative marker genes across clusters where marker genes denote specific populations. **(d)** Box and whisker plots showing the CytoTRACE score distribution of cells within each cluster and organized high to low by mean cluster CytoTRACE score. CytoTRACE estimates the differentiation status of cells (high score = less differentiated; low score = more differentiated). The center line represents the median, box limits are the first and third quartiles, whiskers extend to 1.5 times the interquartile range, and the points beyond the whiskers represent outliers. **(e)** Overlayed UMAP and dot plot of *H2b-egfp* gene expression across identified clusters. **(f)** Unbiased RNA velocity trajectory analysis, split by diet condition. UMAP is overlayed with arrows demonstrating predicted cell trajectories. Cells colored by GFP^+^ subpopulations.

Given that we can detect cells expressing low levels of GFP protein (GFP^lo^) by flow cytometry which we had FACS-sorted in addition to GFP^hi^ cells from *Pdgfrα^H2B-eGFP^* mammary cells for scRNA-seq, we interrogated our single-cell data for expression of the *Gfp* transcript. We generated a sequence of the 3’ end of the *Pdgfrα H2b-egfp* locus, and assigned a gene id to this *Gfp* sequence during alignment. Following data preprocessing and analysis, we found that *Gfp* transcripts were confined to MAPs and not considerably expressed in other clusters **(Fig. 3e and Supplementary Fig. 3b)**, supporting the notion that GFP^lo^ cells which constitute basal cells, luminal progenitors, endothelial and immune cells are descendants of the GFP^hi^ *Pdgfrα*-expressing MAP lineage.

To further validate MAP differentiation potential, we used RNA velocity which is an unbiased pseudotime trajectory analysis method based on unspliced to spliced transcript proportion^21,22^. The analysis unveiled MAP4 as the MAP progenitor cluster giving rise to both MAPs and non-MAP cells **(Fig. 3f)**. In probing distinctions between obese HFD- and lean CD-fed mice, robust trajectories were found to originate from the MAP4 cluster leading into the immune cluster in obesity **(Fig. 3f)**. Upon further characterization, the immune cluster comprised macrophages expressing *Adgre1*, *Cd14*, *Cebpb* and *Cd86*, mature B-cells expressing *Ighm*, and peripheral *Cd3e*-expressing double-negative T-cells **(Supplementary Fig. 4a-c)**. Immune cell proportion differences between CD and HFD indicate that obesity propels the expansion of immunosuppressive macrophages distinguished by *Cd163* and *Hmox1* **(Supplementary Fig. 4d)**. Moreover, trajectories from MAP4 to the basal epithelial and LP1 clusters intensify in obesity, and altered endothelial differentiation is also observed between CD and HFD conditions **(Fig. 3f)**. The basal population in the mammary gland is known to be enriched in bipotent mammary stem cells^23,24^, and stem cells and luminal progenitors are putative cells of origin in breast cancer^25,26^ Together, these orthogonal and unbiased single-cell analyses pinpoint the diverse cell fates and plasticity of MAPs, and the impact of an obesity-inducing diet on augmenting differentiation of MAPs into macrophages, basal cells and luminal progenitors.

### MAPs adopt an invasive cell fate in obesogenic conditions

To define the fate of MAPs in obesity, GFP labeling of MAPs in *Pdgfrα^CreERT^ R26^mTmG^* lineage tracing mice was performed with tamoxifen-mediated Cre induction as previously reported^17^ prior to feeding with HFD or CD **(Fig. 4a)**. The GFP label in *Pdgfrα^CreERT^R26^mTmG^*lineage tracing mice upon induction is permanent, thus allowing us to map the fate of MAPs and progeny in obesogenic conditions. Following the HFD regimen, lineage tracing mice gained significant weight, and showed increased serum insulin, glucose levels and more mammary gland mass **(Fig. 4b-e).** In isolated mammary tissues from lineage tracing mice, GFP-labeled MAPs and progeny were prominent in the periductal space of the HFD group **(Fig. 4f)**. Notably, GFP^+^ cells were observed to make close contacts with EpCAM^+^ epithelial cells, and invade the epithelial compartment in HFD compared to CD. GFP^+^ progeny in the epithelium of obese mice were found to be positive for the basal marker Keratin 14 **(Fig. 4f)**, concordant with the reinforced trajectory of MAPs to basal cells in obesity found in our scRNA-seq data. We also detected GFP^+^ cells that expressed the immune marker CD45 in HFD but not CD, supporting the transition of MAPs into immune cells inferred by our trajectory analysis. Given the evident invasion of GFP^+^ MAP progeny into the mammary gland during obesity, we assessed the state of the basement membrane which typically serves as a barrier between the epithelium and the stroma. Laminins form a critical basement membrane constituent in the mammary gland, and upon immunostaining mammary tissue sections for laminin, we observed a distinct deficiency of intact laminin in mammary tissues of obese HFD mice compared to lean controls **(Fig. 4f)**. This data suggest that MAP dynamics are altered in obesity such that they adopt an invasive cellular state conferring increased access into the epithelium in the absence of a sufficient basement membrane whilst generating increased immune and epithelial progeny.

**Figure 4.**
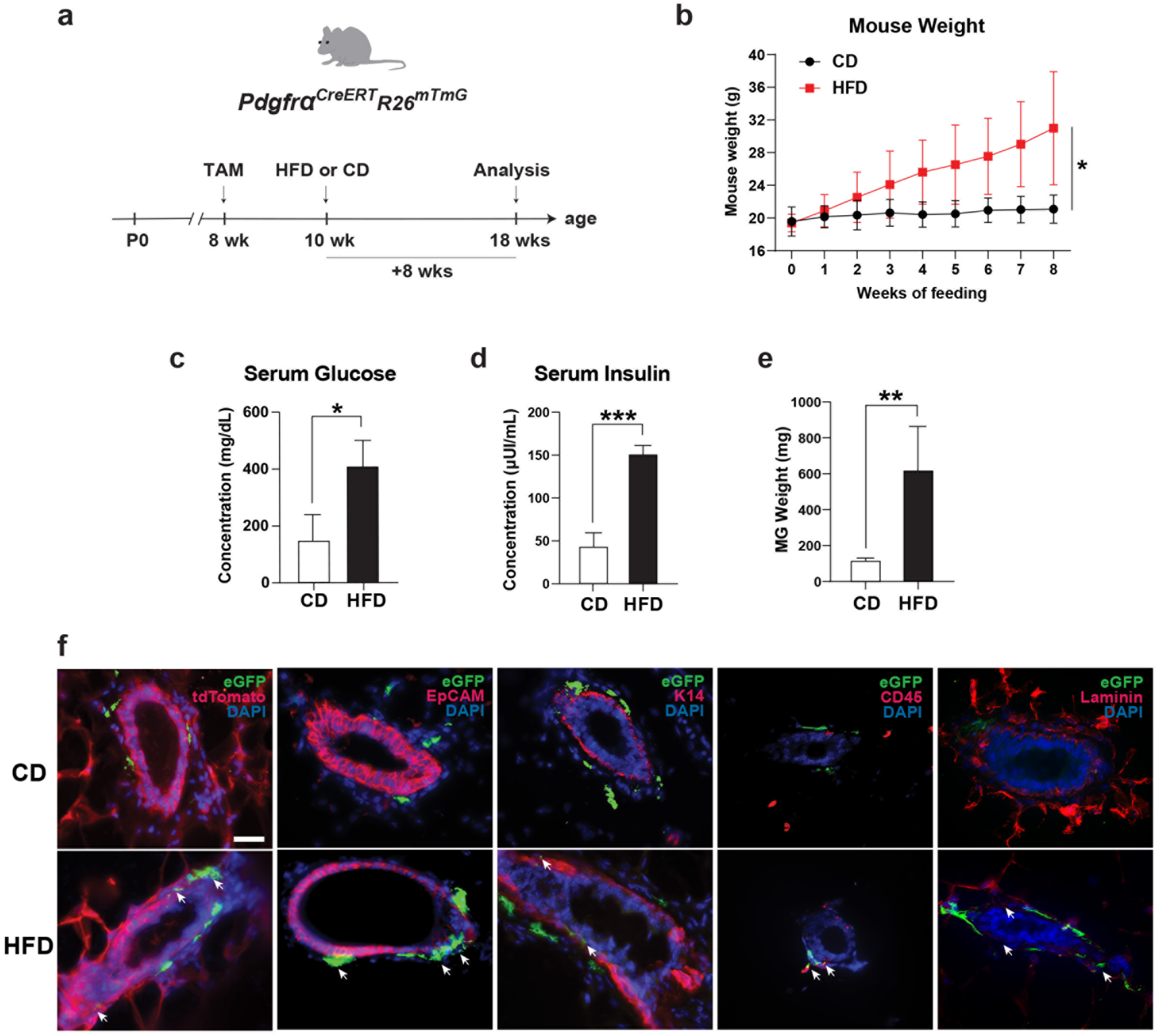
MAPs in obesity diverge into immune and epithelial cells invading the mammary epithelial compartment. **(a)** An inducible *Pdgfrα^CreERT^ R26^mTmG^* lineage tracing model was injected with tamoxifen (TAM) to induce Cre-mediated recombination and permanent GFP labeling of MAPs following which mice were subjected to a HFD or CD regimen. **(b)** Weight gain of HFD-fed lineage tracing mice compared to CD-fed mice over 8 weeks (n=7 for HFD and n=4 for CD). **(c)** Serum insulin, **(d)** Serum glucose (n=3 per group) and **(e)** Mammary gland mass of HFD versus CD mice (n=7 for HFD and n=4 for CD). **(f)** Immunofluorescence on HFD and CD mammary tissue sections for GFP-labeled MAPs and progeny co-stained with the epithelial marker EpCAM, basal marker Keratin 14 (K14), immune marker CD45 and basement membrane component laminin. White arrows indicate GFP progeny or the loss of laminin.

### High-fat diet augments EGFR-mediated responses in MAPs

We then sought to determine mechanistic changes in MAPs resulting from HFD by exploring differentially expressed genes (DEGs; adjusted p-value<0.05) in the MAP4 cluster of our single-cell data **(Supplementary Table 1)** which manifested as the origin of trajectories to immune and epithelial lineages. KEGG pathway analysis^27,28^ performed on MAP4 DEGs between HFD and CD showed an enrichment in cancer and inflammatory pathways including Pathways in cancer, Transcriptional misregulation in cancer, Coronavirus disease, NF-kappa B signaling pathway and MAPK signaling pathway **(Fig. 5a).** Among the top 10 KEGG pathways, *Egfr* and *Mmp3* repeatedly emerged as contributing DEGs **(Fig. 5a and Supplementary Table 2)**. On deeper analysis, both *Egfr* and *Mmp3* were found to be elevated in expression in HFD conditions **(Fig. 5b)**, as indicated by positive log2 fold change values **(Supplementary Table 1)**. Increased *Mmp3* levels may result in greater basement membrane degradation, thus accounting for the lack of intact laminin in HFD mammary glands analyzed in our lineage tracing studies. Furthermore, given its known functions in cell growth and development^29^, Epidermal Growth Factor Receptor (EGFR) is a plausible mechanistic candidate for modulating the proliferation and differentiation from MAPs to non-MAP identities in obesity.

**Figure 5.**
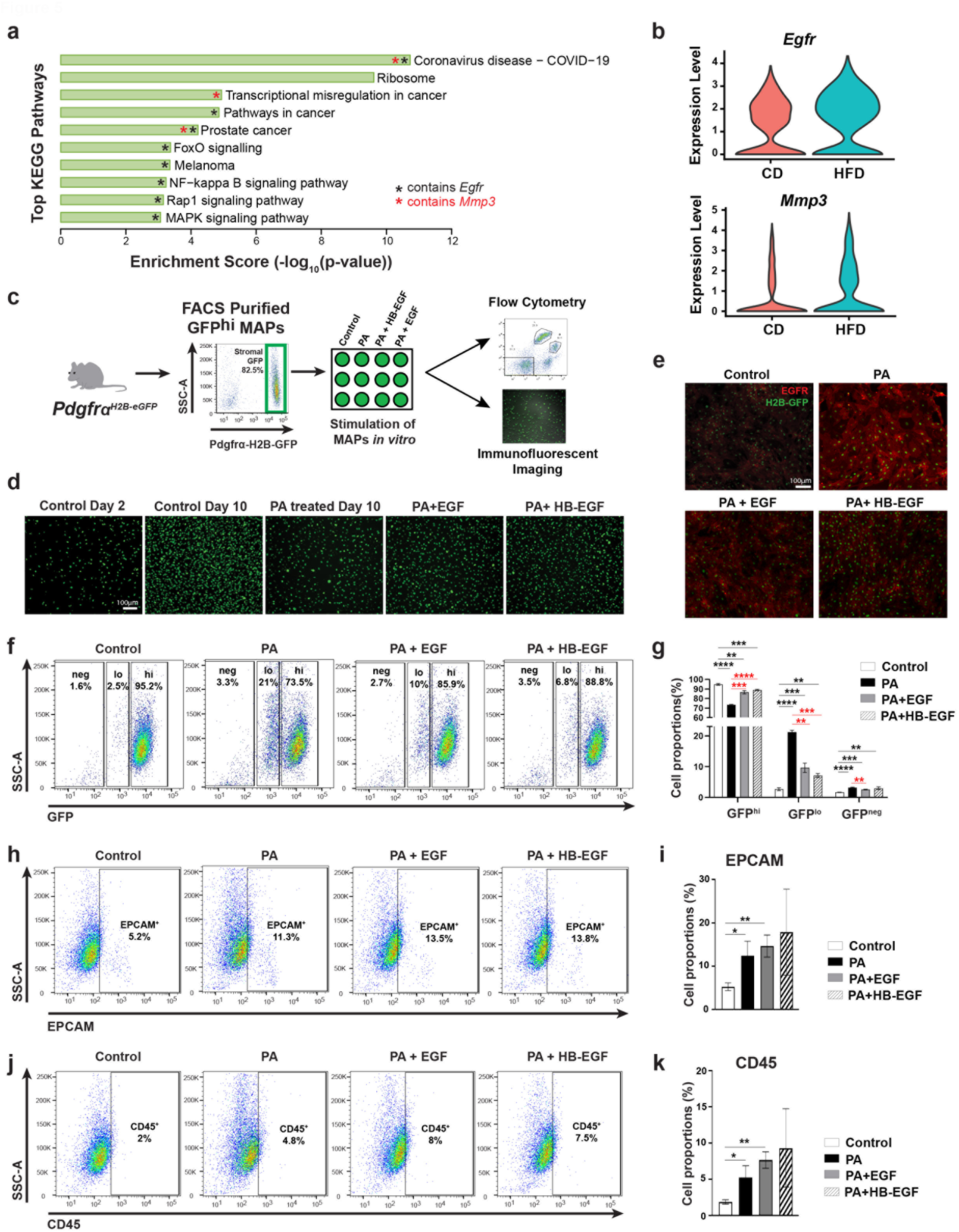
MAPs upregulate EGFR in obesity to propel MAP proliferation, differentiation and plasticity. **(a)** Top 10 most significant KEGG pathways enriched for differentially expressed genes (adjusted p-value <0.05) in MAP4 cluster between obese (HFD) and lean (CD) mice. **(b)** Violin plots illustrating *Egfr* and *Mmp3* gene expression in CD and HFD MAP4 cell population. **(c)** GFP^hi^ MAPs were FACS-purified from *Pdgfrα^H2B-eGFP^*mammary glands and cultured *in vitro* with HFD constituent palmitic acid (PA) in the presence or absence of EGFR ligands EGF or HB-EGF. **(d)** Representative images of endogenous GFP fluorescence of cultured GFP^hi^ MAPs on day 2 after initiation of culture, and on day 10 following PA treatment with or without EGFR ligands (n=3 replicates per condition). **(e)** Immunofluorescence on day 10 MAP cultures for EGFR and H2B-GFP after indicated treatments. **(f)** Representative flow cytometry plots of GFP^hi^, GFP^lo^ and GFP-negative (GFP^neg^) subsets stemming from GFP^hi^ MAPs in control and following culture with PA with or without EGFR stimulation (n=3 replicates per condition). **(g)** Enumeration of GFP^hi^, GFP^lo^ and GFP^neg^ subsets show in (f). Representative flow cytometry plots and quantification for EpCAM^+^ epithelial cells **(h, i)** and CD45^+^ immune cells **(j, k)** generated from MAPs following culture in defined conditions (n=3 replicates per condition). Data represent mean ± s.d. *P< 0.05, **P<0.01, ***P<0.001, ****P<0.0001.

To interrogate sensitivity of MAPs to EGFR-driven responses, we FACS-sorted exclusively GFP^hi^ MAPs with high purity from *Pdgfrα^H2B-eGFP^*mammary glands, and cultured them *in vitro* in the absence (control) or presence of a major high-fat diet component, palmitic acid (PA), with or without additional stimulation by EGFR ligands, EGF or HB-EGF **(Fig. 5c)**. GFP^+^ cells were observed through fluorescence microscopy early on in culture and found to increase in confluency over several days **(Fig. 5d)**. In particular, GFP^hi^ cells appeared to persist in control wells, while GFP^lo^ cells were more apparent in PA and PA+EGFR ligand wells. Immunofluorescent staining for EGFR in these cultures indicated increased EGFR expression especially following PA but also in the presence of EGFR ligands **(Fig. 5e)**. We then performed flow cytometry analysis on the cultures to quantify GFP^hi^ versus GFP^lo^ and GFP negative (GFP^neg^) cells **(Fig. 5f, g)**. Compared to control, treatment of GFP^hi^ cells with PA induced the greatest differentiation into GFP^lo^ and GFP^neg^ cells, whereas EGFR stimulation with ligands in addition to PA led to both significant differentiation into GFP^lo^ and GFP^neg^ cells compared to control, and more proliferation of GFP^hi^ cells compared to treatment with PA alone. When analyzed for the expression of the epithelial marker EpCAM and immune marker CD45, we were able to detect an increase in EpCAM^+^ cells **(Fig. 5h, i)** and CD45^+^ cells **(Fig. 5j, k)** in these MAP cultures following PA and EGFR stimulation compared to controls. Our results demonstrate a marked sensitivity and plasticity of MAPs to high-fat diet and EGFR activation, impelling their proliferation and differentiation into non-MAP cell types. Thus, obesogenic conditions resulting in excess adiposity elicit dynamic EGFR-mediated MAP activity in the mammary gland to ensure a divergent MAP pool that predominantly evolves into macrophages and epithelial progenitor cells, culminating in an inflammatory, cancer-susceptible tissue state **(Fig. 6)**.

**Figure 6.**
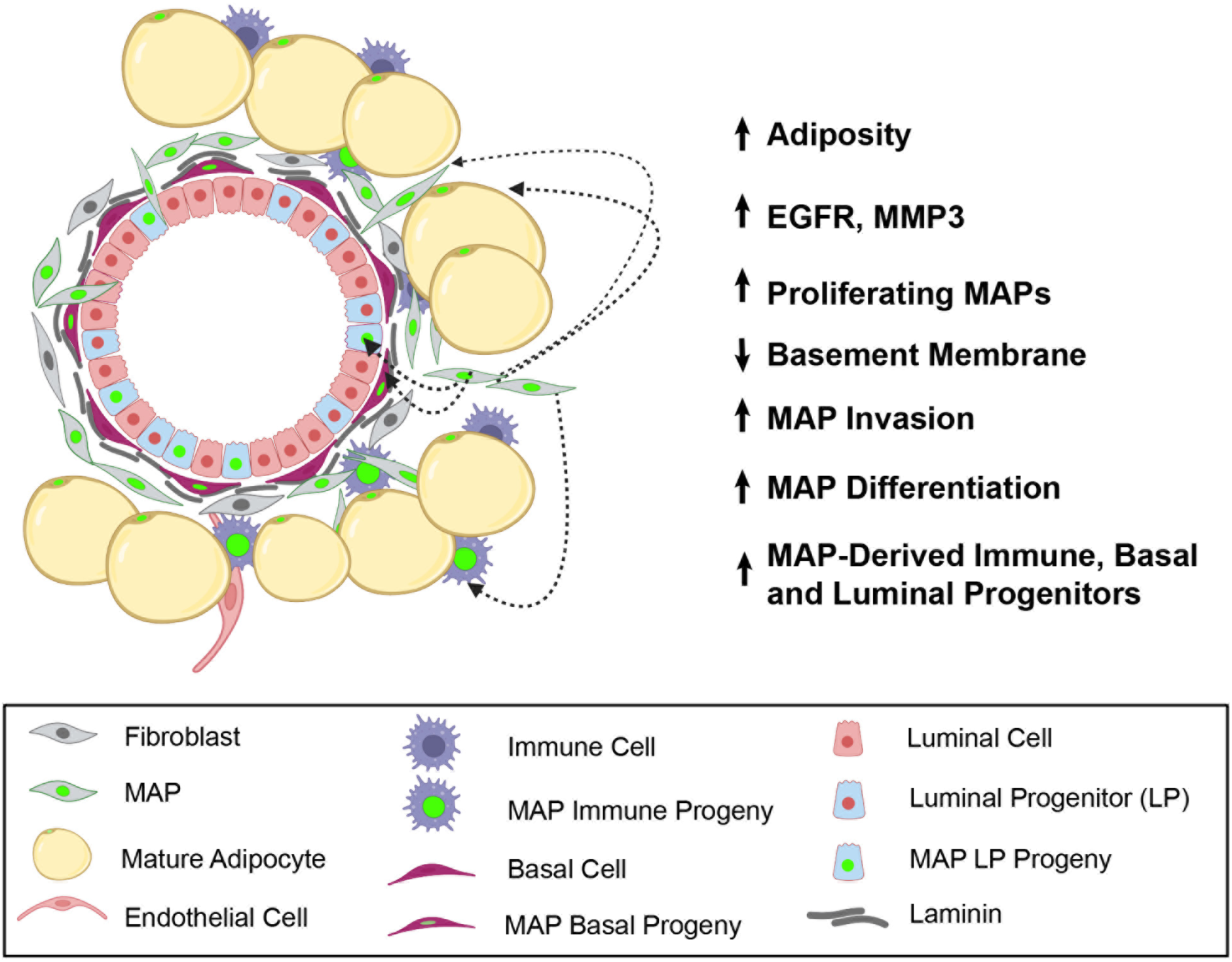
Conceptual model of MAP dynamics in the mammary milieu during excess adiposity. Mammary adipocyte progenitors (MAPs) are activated in obesogenic conditions through enhanced EGFR expression and sensitivity that promote MAP proliferation and differentiation towards diverse cell fates that include immune cells, luminal progenitors and basal epithelial cells. Loss of a laminin-rich basement membrane likely due to increased *Mmp3* facilitates MAP invasion into the mammary epithelial space. These dynamics of MAPs converge to fuel a heightened cancer-prone mammary tissue state in obesity.

## Discussion

Adipocyte progenitors in adipose tissue depots are crucial for ensuring adipocyte differentiation, but uncontrolled fat accretion in obesity thwarts their adipogenic capacity along with the formation of a fibrotic, inflammatory adipose tissue niche ^14,15^. Mammary stromal tissue is home to abundant adipocytes, and we have previously reported the presence of PDGFRα^+^ adipocyte progenitors adjacent to the mammary epithelial parenchyma, which we refer to as MAPs, that have adipogenic potential^17^. MAPs also have an astonishing property to generate epithelial cells during expansion of the mammary epithelium in pregnancy or following estrogen and progesterone sex hormone stimulation. Obesity is recognized to increase breast cancer risk^2–6^, but the state and contribution of local tissue MAPs to altered mammary gland homeostasis in obesity remained elusive until now.

In this study, we elucidate amplified PDGFRα^+^ MAPs in the mammary microenvironment following an obesogenic HFD regimen in C57BL/6J mice. This is a well-established obesity model which is known to mimic diet-induced obesity and associated metabolic abnormalities in humans^30,31^. We find that PDGFRα cell surface protein, *Pdgfrα* transcript and *H2b-Gfp* transcript in *Pdgfrα* reporter mice remain specific to stromal MAPs in obesity as seen by immunostaining and scRNA-seq analysis. Thus, we show that flow cytometry detection of low intensity GFP protein in non-MAP mammary tissue cell types in fluorescent reporter mice of both CD and HFD groups indicates their derivation from the MAP lineage. Through unsupervised scRNA-seq analysis, we also illustrate that MAPs have higher developmental potential based on gene count compared to non-MAP cells namely luminal progenitors, basal, immune and endothelial cells in mammary tissue that retain very low levels of the GFP reporter protein. In obesity, we find a dramatic boost in the cell fate trajectory of MAPs to macrophages, basal epithelial cells and a luminal progenitor subset. Trajectory inference analysis is supported by lineage tracing data that show MAPs transition into epithelial cells and immune progeny in obesogenic conditions. This plasticity of MAPs accompanies an invasive cell state which enables MAPs to cross into the epithelial niche, facilitated by a loss of basement membrane integrity.

Obesity has been shown to increase the activity of epithelial progenitors in the mouse and human mammary gland^32^ which are immature cells and candidate cells of origin in breast cancer^33^. Our results identify elevated MAPs as a source of increased epithelial progenitors in obesity. Also, immune cells, especially macrophages, and their interactions with adipocytes and other cells in mammary tissue are implicated in promoting low-grade chronic inflammation in breast adipose tissue and contributing to carcinogenesis^34^. In the immune population emanating from MAPs, we show that MAPs preferentially commit to a macrophage cell fate trajectory in obesity. Hence, dysregulated MAPs present a cellular origin for diverse cell types that shape a cancer-prone mammary gland in obesity. Intriguingly, mature adipocytes are reported to de-differentiate into mesenchymal adipocyte progenitor-like cells in mammary cancer and this de-differentiation process is associated with a pro-tumorigenic microenvironment^35^. Thus, a tissue niche dominant in adipocyte progenitors is likely to be also favorable for mammary cancer progression independent of obesity.

Mechanistic links between obesity and breast cancer risk have been attributed mostly to altered levels of insulin, IGF-1, adipokines, estrogen and inflammatory mediators^1^. These secreted factors are elevated in adipose tissue through a poorly understood interplay of cell types including adipocytes, immune cells, endothelial cells, a heterogeneous cell population referred to as adipose-derived stromal cells, and the extracellular matrix^36^. We show here that increased MAPs in mammary tissue during obesity have heightened activation of inflammatory and cancer-related pathways in which *Egfr* levels are elevated. Indeed, MAPs expressed greater levels of EGFR, and demonstrated increased proliferation and differentiation into cell types expressing immune and epithelial markers upon exposure to high-fat diet conditions and EGFR stimulation. Collectively, findings from our research illuminate the preponderance of primitive adipocyte progenitors in the mammary gland during obesity which become rewired to generate cell lineages pertinent to cancer development. Thus, targeting MAPs may provide a new avenue for controlling the deviant microenvironment in obesity to reduce mammary cancer risk.

## Methods

### Mice, diets and lineage tracing

All mice were on the C57BL/6J background and maintained on a 12 h light, 12 h dark cycle with access to food and water ad libitum. High-fat diet (HFD; F3282) and matched control diet (CD; F4031) were purchased from Bio-Serv. C57BL/6J wild-type (Stock# 000664), *Pdgfrα^H2B-eGFP^* (Stock# 007669), *Pdgfrα^CreERT^* (Stock# 018280) and *Rosa26^mTmG^* (Stock# 007676) strains were procured from the Jackson Laboratory. All female mice analyzed were validated to be in a non-diestrus phase of the estrous cycle by vaginal cytology given that diestrus is marked by significant cellular and molecular changes in the mammary gland^37^. Inducible lineage tracing in the *Pdgfrα^CreERT^R26^mTmG^*model was performed in adult female mice following a single intraperitoneal injection of 1.25 mg of 4-hyroxytamoxifen (TAM; Sigma) diluted in sunflower seed oil per 25 g body weight to induce Cre-mediated recombination and GFP labeling. Treatment and care of wild-type and transgenic murine models were in accordance with protocols reviewed by the Institutional Animal Care and Use Committee (IACUC) at the University of Texas at Dallas.

### Serum assays

Blood was collected in serum collection tubes (Fisher Scientific). Serum measurements for insulin and glucose were determined using the glucose colorimetric detection kit (Invitrogen) and the mouse insulin solid-phase sandwich ELISA kit (Invitrogen) according to instructions provided by the manufacturer.

### Mammary tissue whole mounts, immunostaining and imaging

Isolated inguinal mammary glands were spread on a glass slide and fixed overnight in Carnoy’s fixative at room temperature. Mounted glands were subsequently rehydrated in descending concentrations of ethanol (70 to 10%) for 15 mins and then distilled water followed by staining overnight with carmine alum. Glands were then washed in ascending concentrations of ethanol (70 to 100%), cleared in xylene and mounted with Permount medium and a glass cover slip. Whole mounts were imaged on an Olympus SZ61 stereo microscope. For immunofluorescence, freshly dissected mammary glands were fixed in 4% paraformaldehyde at room temperature, washed with PBS, incubated in 30% sucrose overnight at 4°C and embedded in OCT after which samples were stored in −80°C and cryosectioned at 5-10 μm thickness for immunofluorescence. Cryosections were incubated in blocking buffer (5% normal donkey serum/1% BSA/0.2% Triton in PBS) for 1 h at room temperature and then in primary antibodies diluted in blocking buffer without Triton overnight at 4°C. Sections were then washed with PBS and incubated with fluorophore-conjugated secondary antibodies for 1 h at room temperature, washed and mounted using ProLong Gold Anti-fade reagent with or without DAPI nuclear stain. For immunostaining cell cultures, cells were fixed with 4% paraformaldehyde for 20 mins, washed with PBS, permeabilized with 0.1% Triton and incubated in blocking buffer with 1% BSA, 5% normal donkey serum in PBS followed by incubation with primary antibodies in blocking buffer overnight. The next day, cells were washed with PBS and incubated with fluorophore-conjugated secondary antibodies diluted in blocking buffer for 1 h at room temperature, washed with PBS and imaged.

Primary antibodies used: anti-PDGFRα (R&D systems, Cat.# AF1062), anti-EpCAM (Abcam, Cat.# ab71916), anti-GFP (Abcam, Cat.# ab13970), anti-Ki67 (eBioscience, Cat.#14-5698-82), anti-tdTomato (Takara, Cat.# 632496), anti-K14 (Biolegend, Cat.# 905301), anti-laminin (Abcam, Cat.# ab11575), anti-CD45 (Cell signaling, Cat.# 70257) and anti-EGFR (Sigma, Cat.# E1282). Secondary antibodies used: anti-goat, anti-rabbit, anti-chicken, anti-rat antibodies conjugated to AlexaFluor 647, AlexaFluor Cy3, AlexaFluor 488 or Dylight 405 (Jackson ImmunoResearch). Immunostained sections were imaged on an Echo Revolve microscope in the upright setting using a 40X Plan Fluorite with Phase 0.75 NA objective lens, and at least 3-5 fields were imaged per tissue section. Cell culture immunofluorescence was imaged on the inverted Revolve setting using 10X Plan Fluorite with Phase 0.3 NA objective lens. Image acquisition was performed with the iOS-based Echo Pro app interface.

### Tissue dissociation, flow cytometry and FACS

Mammary glands were dissociated into single cells as previously described^17,37^. Briefly, glands were digested with 750Uml^-1^ collagenase and 250Uml^-1^ hyaluronidase in DMEM: F12 media for 2.5 hours at 37°C. The digestion mixture with tissues was then vortexed, washed in Hank’s balanced salt solution with 2% FBS followed by red blood cell lysis in ammonium chloride solution, incubation in 0.25% trypsin for 2 min and 5 mgml^-1^ dispase with 0.1 mgml^-1^ DNaseI for 2 min, and then filtered through a 40μm mesh to generate single cells. All dissociation reagents were obtained from STEMCELL Technologies. For flow cytometry analysis, cells were labeled with Pe-Cy7-conjugated antibodies to CD45 (Ebioscience; Cat.# 25-0451), CD31 (Ebioscience, Cat.# 25-0311) and Ter119 (Ebioscience, Cat.# 25-5921) for immune, endothelial and erythrocyte populations respectively which were jointly referred to as lineage^+^ (Lin^+^) cells, and excluded from analysis of mammary epithelial and stromal cells. The following antibodies were used to identify mammary epithelial subpopulations and stromal cells enriched in MAPs: anti-CD49f-APC (R&D, Cat.# FAB13501A), anti-EpCAM-APC/Cy7 (Biolegend, Cat.# 118217), anti-CD140 (PDGFRα)-PE (Biolegend, Cat.# 135905) or anti-CD140 (PDGFRα)-BV421 (Biolegend, Cat.# 135923). Dead cells were excluded with Zombie UV dye (Biolegend, Cat.# 423107). Flow cytometry was performed using a Fortessa cell analyzer with FACsDiva software (BD), and FlowJo software (Tree Star, Inc.) was used for downstream analysis. Cell sorting by FACS was done on a FACSAria Fusion (BD) and the purity of sorted populations was routinely >98%.

### *In vitro* MAP stimulation

FACS-sorted GFP^hi^ MAPs from mammary glands of *Pdgfrα^H2B-eGFP^* mice were seeded onto collagen-coated wells of a 24-well plate at 20,000 cells per well and cultured in DMEM media with 10% FBS and 10ng/ml bFGF (STEMCELL Technologies). Following 5 days of culture, Palmitic Acid (30μM; Cayman Chemical) was added alone or with EGF (20 ng/ml; STEMCELL Technologies) or HB-EGF (20 ng/ml; Sigma) in DMEM/10% FBS. Control continued to be cultured in media with bFGF alone. Cultures were maintained for 5 days with treatments and cells were subsequently analyzed by immunofluorescence and flow cytometry.

### *Pdgfrα H2b-eGfp* 3’ terminal sequencing

For sequencing the 3’ end of the *Pdgfrα H2b-eGfp* locus, tissue was taken from a heterozygous *Pdgfrα^H2B-eGFP^* mouse and incubated overnight in digestion buffer (1M Tris-HCl, 0.25M EDTA, 10% SDS, 0.4mg/mL Proteinase K, pH 8.0). Primers were designed based on the genetic components stated to be included in the design of the original construct^38,39^. The sample was then diluted nine-fold with distilled water and the 3’ end of the *H2b-eGfp* locus was amplified using PCR, run on a 1.5% agarose gel, and isolated using a PureLink Quick Gel Extraction Kit. Isolated DNA was then mixed with one of two sequencing primer and sent to the UT Dallas Genome Center for Sanger sequencing using an Applied Biosystems SeqStudio Genetic analyzer. Resulting sequence fragments were aligned using the NIH Nucleotide BLAST program to generate the composite sequence of *H2b-eGfp* 3’ end. The resulting sequence was further processed by excluding variable nucleotide results and the remaining 676bp sequence submitted for single cell RNA sequencing alignment. Primers used for both amplification and sequencing were WT-F: CCCTTGTGGTCATGCCAAAC and H2B-Amp-F: GGCATGGACGAGCTGTACAAG.

### Single-cell RNA-sequencing (scRNA-seq)

Live GFP^+^ cells were FACS-sorted from *Pdgfrα^H2B-eGFP^*mice fed a HFD and matched CD for scRNA-seq. Standard procedures provided from 10X Genomics were used for sample preparation and Chromium Next GEM single cell 3’ gene expression library construction. Sequencing was conducted on an Illumina Nextseq 2000 P2 flow cell 100bp and sequencing read was set up as follows: Read1:28 cycles, I1: 10 cycles, Index 2: 10 cycles, Read2: 90 cycles. Sample demultiplexing, alignment, and filtering were performed with Cell Ranger software (v5.0.1) with the mm10 mouse genome as reference. *H2b-gfp* gene sequence was annotated under gene name *“GFP_new_id”.* For CD, 8,219 cells were recovered with a median UMI count of 3,602 per cell, a mean reads per cell of 24,405, and a median genes per cell of 1,629. For HFD, 8,612 cells were recovered with a median UMI count of 3,708 per cell, a mean reads per cell of 23,224, and a median genes per cell of 1,648. Genes *Gm42418* and *AY036118* were removed, as these overlap with the rRNA element Rn45s and represent rRNA contamination. Homotypic and heterotypic doublets were filtered with DoubletFinder (v2.0.4). In CD, 7,679 cells were classified as singlets at a doublet frequency of 6.57%. In HFD, 8,019 cells were classified as singlets at a doublet frequency of 6.89%.

### scRNA-seq analysis

Clustering and gene expression were visualized with the Seurat package (v4.4.0) in RStudio. Unsupervised clustering with a resolution of 0.4 identified eleven clusters, and were annotated based on validated gene expression profiles. RNA Velocity Unbiased Trajectory Inference Analysis based on spliced and unspliced transcript proportions was performed with velocyto (v0.17) and ScVelo PyPi (v0.2.5) in Python (v3.11.4). Unsupervised hierarchical differentiation states and stemness were predicted with CytoTRACE (v0.3.3), with a higher CytoTRACE score inferring a more progenitor-like population. KEGG Pathway enrichment analyses were conducted with differentially expressed genes (FDR<0.05) derived from Seurat FindMarkers function using the NIH DAVID Knowledgebase (v2023q3).

### Statistical Analyses

For *in vivo* experiments, n represents a distinct biological replicate and data for biological replicates were pooled from independent experiments. For *in vitro* experiments, n represents an individual technical replicate. Experiments were repeated at least two independent times with successful replication. Data were analyzed using GraphPad prism software and reported as mean ± standard error of the mean (s.e.m). Comparison of data between two groups was made using Student’s *t*-test (two-tailed). Statistical significance is recognized at *p*<0.05.

## Data availability

Processed and raw UMI count matrices will be deposited in the NCBI Gene Expression Omnibus. Code will be made available on GitHub.

## Acknowledgements

This work was supported by a Victoria’s Secret Global Fund for Women’s Cancers 2023 Career Development Award, in partnership with Pelotonia & AACR, Grant Number 23-20-73-JOSH and UT Dallas Startup funds to PAJ. SK holds a Graduate Research and Cancer Education Fellowship from UT Dallas. The authors thank Philipp Scherer (UT Southwestern) for high-fat diet recommendations and provisions. The authors acknowledge the support of Yeunhee Kim from the UT Dallas Genome Center for library preparation and sequencing, UT Dallas Cyberinfrastructure & Research Services Department for providing access to high-performance computing, Jacob Henderson from the UT Dallas FACS Core for assistance with cell sorting, and the UT Southwestern Bioinformatics Core for data preprocessing in Cell Ranger.

## Author contributions

PAJ conceived, designed and directed the study. SAK contributed to study design and performed majority of the experiments and biological data analyses. PN executed all scRNA-seq analyses. JPSS contributed to immunostaining and imaging of cells and tissues. SM carried out flow cytometry and associated data analyses for cell culture experiments. DN assisted with tissue processing and made cryosections for all tissues analyzed in the study. DGS generated *the Pdgfrα H2b-Gfp* 3’ terminal sequence. ZX provided guidance and recommendations for scRNA-seq analyses. SAK, PN, JPSS and SM contributed to figure compilation. PAJ and SK acquired funds for the study. PAJ, PN and SK wrote the manuscript.

## References

1 Avgerinos, K. I., Spyrou, N., Mantzoros, C. S. & Dalamaga, M. Obesity and cancer risk: Emerging biological mechanisms and perspectives. Metabolism 92, 121–135 (2019). 10.1016/j.metabol.2018.11.001

2 Picon-Ruiz, M., Morata-Tarifa, C., Valle-Goffin, J. J., Friedman, E. R. & Slingerland, J. M. Obesity and adverse breast cancer risk and outcome: Mechanistic insights and strategies for intervention. CA Cancer J Clin 67, 378–397 (2017). 10.3322/caac.21405

3 Kawai, M., Malone, K. E., Tang, M. T. & Li, C. I. Height, body mass index (BMI), BMI change, and the risk of estrogen receptor-positive, HER2-positive, and triple-negative breast cancer among women ages 20 to 44 years. Cancer 120, 1548–1556 (2014). 10.1002/cncr.28601

4 Nagrani, R. et al. Central obesity increases risk of breast cancer irrespective of menopausal and hormonal receptor status in women of South Asian Ethnicity. Eur J Cancer 66, 153–161 (2016). 10.1016/j.ejca.2016.07.022

5 Amadou, A. et al. Overweight, obesity and risk of premenopausal breast cancer according to ethnicity: a systematic review and dose-response meta-analysis. Obes Rev 14, 665–678 (2013). 10.1111/obr.12028

6 Millikan, R. C. et al. Epidemiology of basal-like breast cancer. Breast Cancer Res Treat 109, 123–139 (2008). 10.1007/s10549-007-9632-6

7 Yedjou, C. G. et al. Health and Racial Disparity in Breast Cancer. Adv Exp Med Biol 1152, 31–49 (2019). 10.1007/978-3-030-20301-6_3

8 Stringer-Reasor, E. M., Elkhanany, A., Khoury, K., Simon, M. A. & Newman, L. A. Disparities in Breast Cancer Associated With African American Identity. Am Soc Clin Oncol Educ Book 41, e29–e46 (2021). 10.1200/EDBK_319929

9 Park, J., Morley, T. S., Kim, M., Clegg, D. J. & Scherer, P. E. Obesity and cancer--mechanisms underlying tumour progression and recurrence. Nat Rev Endocrinol 10, 455–465 (2014). 10.1038/nrendo.2014.94

10 Zeve, D., Tang, W. & Graff, J. Fighting fat with fat: the expanding field of adipose stem cells. Cell stem cell 5, 472–481 (2009). 10.1016/j.stem.2009.10.014

11 Lee, Y. H., Petkova, A. P., Mottillo, E. P. & Granneman, J. G. In vivo identification of bipotential adipocyte progenitors recruited by beta3-adrenoceptor activation and high-fat feeding. Cell Metab 15, 480–491 (2012). 10.1016/j.cmet.2012.03.009

12 Berry, R. & Rodeheffer, M. S. Characterization of the adipocyte cellular lineage in vivo. Nat Cell Biol 15, 302–308 (2013). 10.1038/ncb2696

13 Cattaneo, P. et al. Parallel Lineage-Tracing Studies Establish Fibroblasts as the Prevailing In Vivo Adipocyte Progenitor. Cell Rep 30, 571–582 e572 (2020). 10.1016/j.celrep.2019.12.046

14 Ghaben, A. L. & Scherer, P. E. Adipogenesis and metabolic health. Nat Rev Mol Cell Biol 20, 242–258 (2019). 10.1038/s41580-018-0093-z

15 Marcelin, G. et al. A PDGFRalpha-Mediated Switch toward CD9(high) Adipocyte Progenitors Controls Obesity-Induced Adipose Tissue Fibrosis. Cell Metab 25, 673–685 (2017). 10.1016/j.cmet.2017.01.010

16 Han, M. S. et al. A feed-forward regulatory loop in adipose tissue promotes signaling by the hepatokine FGF21. Genes Dev 35, 133–146 (2021). 10.1101/gad.344556.120

17 Joshi, P. A. et al. PDGFRalpha(+) stromal adipocyte progenitors transition into epithelial cells during lobulo-alveologenesis in the murine mammary gland. Nat Commun 10, 1760 (2019). 10.1038/s41467-019-09748-z

18 Beyaz, S. et al. High-fat diet enhances stemness and tumorigenicity of intestinal progenitors. Nature 531, 53–58 (2016). 10.1038/nature17173

19 Rodeheffer, M. S., Birsoy, K. & Friedman, J. M. Identification of white adipocyte progenitor cells in vivo. Cell 135, 240–249 (2008). 10.1016/j.cell.2008.09.036

20 Gulati, G. S. et al. Single-cell transcriptional diversity is a hallmark of developmental potential. Science 367, 405–411 (2020). 10.1126/science.aax0249

21 La Manno, G. et al. RNA velocity of single cells. Nature 560, 494–498 (2018). 10.1038/s41586-018-0414-6

22 Bergen, V., Lange, M., Peidli, S., Wolf, F. A. & Theis, F. J. Generalizing RNA velocity to transient cell states through dynamical modeling. Nat Biotechnol 38, 1408–1414 (2020). 10.1038/s41587-020-0591-3

23 Stingl, J. et al. Purification and unique properties of mammary epithelial stem cells. Nature 439, 993–997 (2006). 10.1038/nature04496

24 Rios, A. C., Fu, N. Y., Lindeman, G. J. & Visvader, J. E. In situ identification of bipotent stem cells in the mammary gland. Nature 506, 322–327 (2014). 10.1038/nature12948

25 Shehata, M. et al. Phenotypic and functional characterisation of the luminal cell hierarchy of the mammary gland. Breast Cancer Res 14, R134 (2012). 10.1186/bcr3334

26 Molyneux, G. et al. BRCA1 basal-like breast cancers originate from luminal epithelial progenitors and not from basal stem cells. Cell Stem Cell 7, 403–417 (2010). 10.1016/j.stem.2010.07.010

27 Kanehisa, M., Furumichi, M., Sato, Y., Kawashima, M. & Ishiguro-Watanabe, M. KEGG for taxonomy-based analysis of pathways and genomes. Nucleic Acids Res 51, D587–D592 (2023). 10.1093/nar/gkac963

28 Kanehisa, M. & Goto, S. KEGG: kyoto encyclopedia of genes and genomes. Nucleic Acids Res 28, 27–30 (2000). 10.1093/nar/28.1.27

29 Sibilia, M. et al. The epidermal growth factor receptor: from development to tumorigenesis. Differentiation 75, 770–787 (2007). 10.1111/j.1432-0436.2007.00238.x

30 Hariri, N. & Thibault, L. High-fat diet-induced obesity in animal models. Nutr Res Rev 23, 270–299 (2010). 10.1017/S0954422410000168

31 Kanasaki, K. & Koya, D. Biology of obesity: lessons from animal models of obesity. J Biomed Biotechnol 2011, 197636 (2011). 10.1155/2011/197636

32 Chamberlin, T., D’Amato, J. V. & Arendt, L. M. Obesity reversibly depletes the basal cell population and enhances mammary epithelial cell estrogen receptor alpha expression and progenitor activity. Breast Cancer Res 19, 128 (2017). 10.1186/s13058-017-0921-7

33 Fu, N. Y., Nolan, E., Lindeman, G. J. & Visvader, J. E. Stem Cells and the Differentiation Hierarchy in Mammary Gland Development. Physiol Rev 100, 489–523 (2020). 10.1152/physrev.00040.2018

34 Iyengar, N. M., Hudis, C. A. & Dannenberg, A. J. Obesity and inflammation: new insights into breast cancer development and progression. Am Soc Clin Oncol Educ Book 33, 46–51 (2013). 10.14694/EdBook_AM.2013.33.46

35 Zhu, Q. et al. Adipocyte mesenchymal transition contributes to mammary tumor progression. Cell Rep 40, 111362 (2022). 10.1016/j.celrep.2022.111362

36 Hillers-Ziemer, L. E., Kuziel, G., Williams, A. E., Moore, B. N. & Arendt, L. M. Breast cancer microenvironment and obesity: challenges for therapy. Cancer Metastasis Rev 41, 627–647 (2022). 10.1007/s10555-022-10031-9

37 Joshi, P. A. et al. Progesterone induces adult mammary stem cell expansion. Nature 465, 803–807 (2010). 10.1038/nature09091

38 Hamilton, T. G., Klinghoffer, R. A., Corrin, P. D. & Soriano, P. Evolutionary divergence of platelet-derived growth factor alpha receptor signaling mechanisms. Mol Cell Biol 23, 4013–4025 (2003). 10.1128/MCB.23.11.4013-4025.2003

39 Kanda, T., Sullivan, K. F. & Wahl, G. M. Histone-GFP fusion protein enables sensitive analysis of chromosome dynamics in living mammalian cells. Curr Biol 8, 377–385 (1998). 10.1016/s0960-9822(98)70156-3

