## Supplementary Data for "Obesity modifies cell fate plasticity of *Pdgfrα-*expressing adipocyte progenitors to promote an aberrant mammary microenvironment"

**Kwende et al.**

Supplementary Data

Supplementary Figure 1

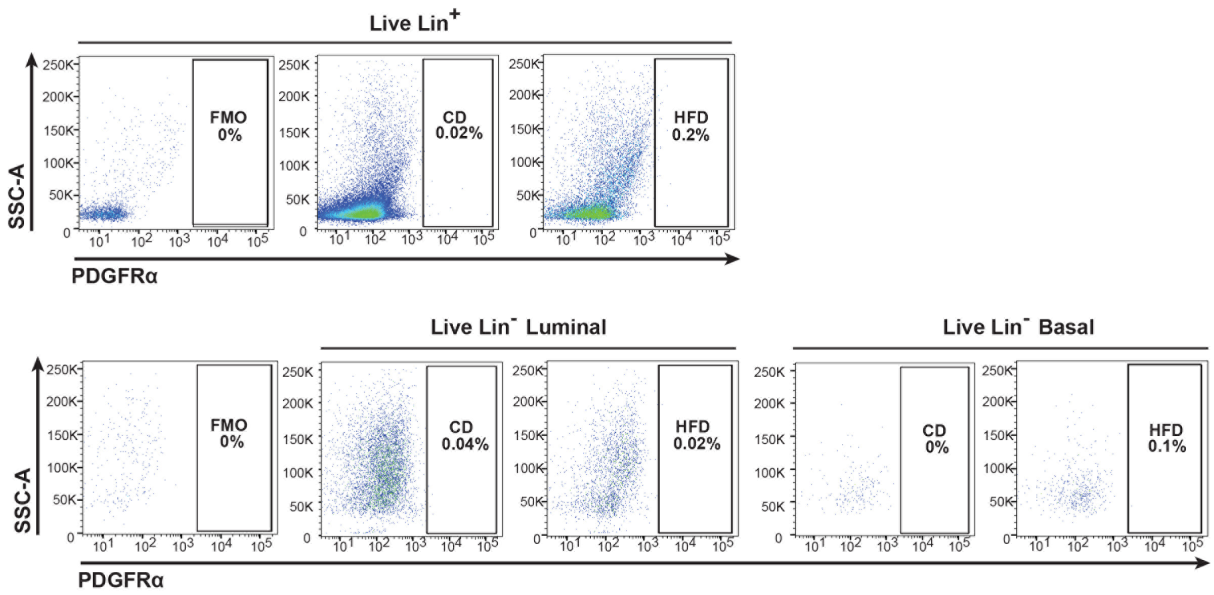


**Supplementary Figure 1. Absence of PDGFRα$^+$ cells in non-MAP-enriched mammary cell populations.** Flow cytometry analysis of PDGFRα$^+$ cells in live lineage$^+$ (CD45$^+$ CD31$^+$ Ter119$^+$, Lin$^+$) population and Lineage$^-$ (Lin$^-$) luminal and basal epithelial subpopulations in mammary glands from mice fed a high-fat diet (HFD) or matched control diet (CD) (n=7 mice per group); Fluorescence Minus One (FMO) control for gating PDGFRα$^+$ cells.

Supplementary Figure 2

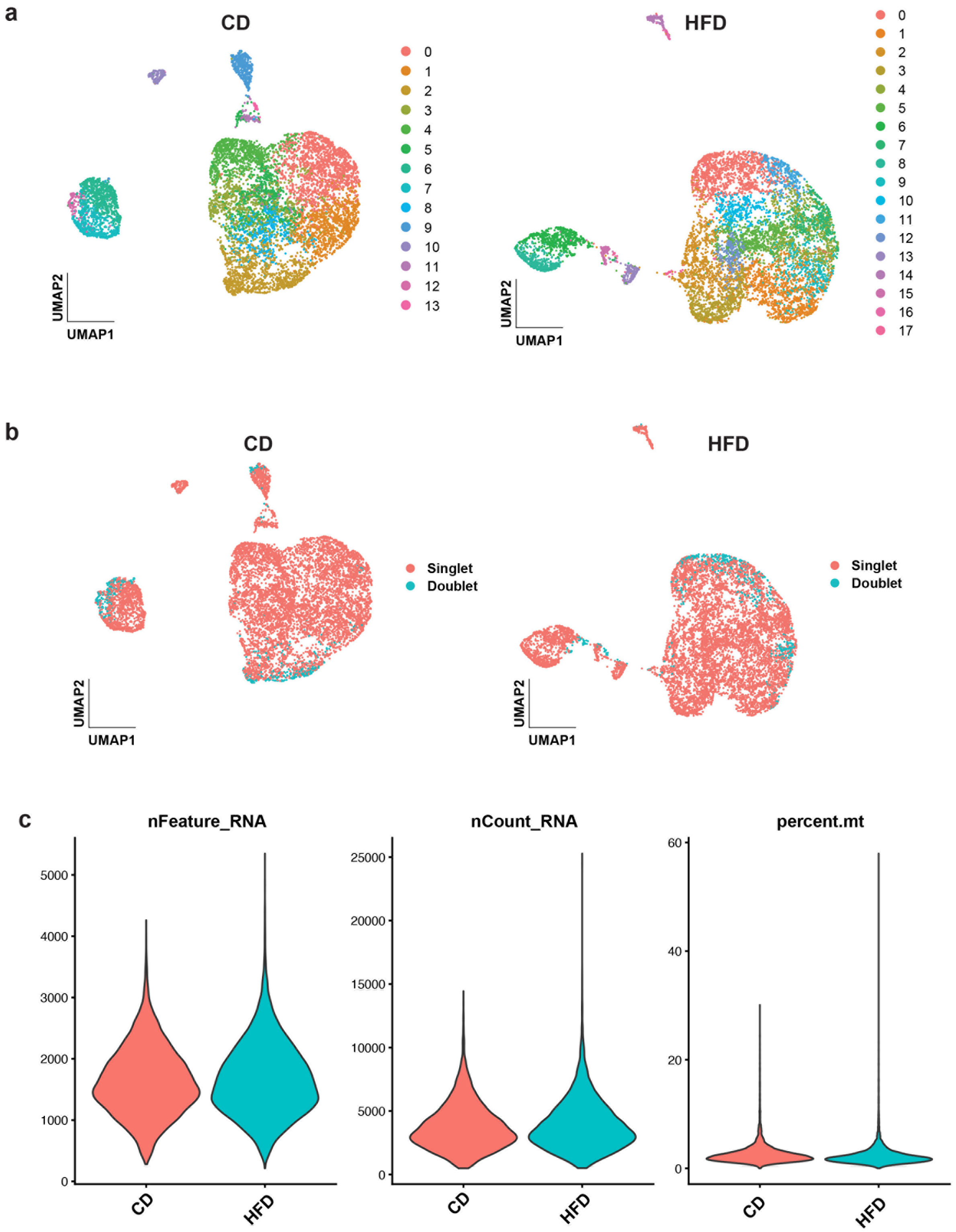


**Supplementary Figure 2. Quality control and preprocessing of single-cell RNA sequencing data.**

**(a)** Clustered UMAPs from Cell Ranger output for HFD and CD samples, after rRNA contamination removal. **(b)** UMAPs of original clustering, showing doublets and singlets. CD had 540 doublets (~6.58%). HFD had 593 doublets (~6.89%). Doublet removal was conducted in concordance with 10x multiplet rates. **(c)** Quality control metrics after rRNA contaminant and doublet removal indicating normal range values for CD and HFD.

Supplementary Figure 3

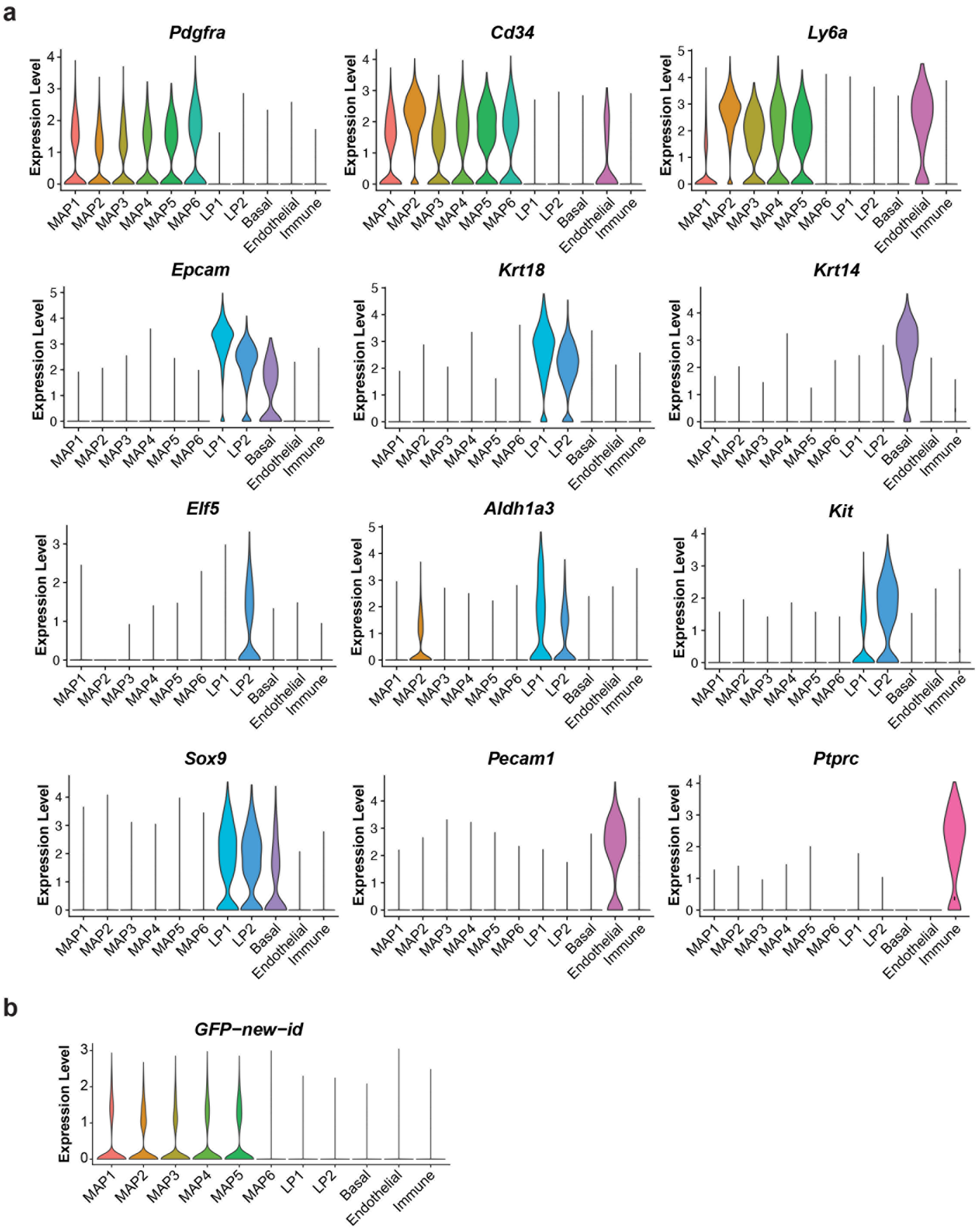


**Supplementary Figure 3. Violin plots of gene expression used to determine cluster identity and *H2b-Gfp*-expressing clusters. (a)** Violin plots of marker genes. *Pdgfr*$\alpha$ expression indicates the presence of 6 MAP clusters. *Epcam* shows the presence of 3 epithelial clusters which is further split into 2 luminal progenitor (LP) clusters by *Ly6a*, *Krt18*, *Elf5*, *Aldh1a3*, *Kit and Sox9*, and 1 basal cluster by *Krt14*. *Pecam1* marks an endothelial cluster whereas *Ptprc* marks an immune cluster. **(b)** *H2b-Gfp* expression is restricted to MAPs.

Supplementary Figure 4

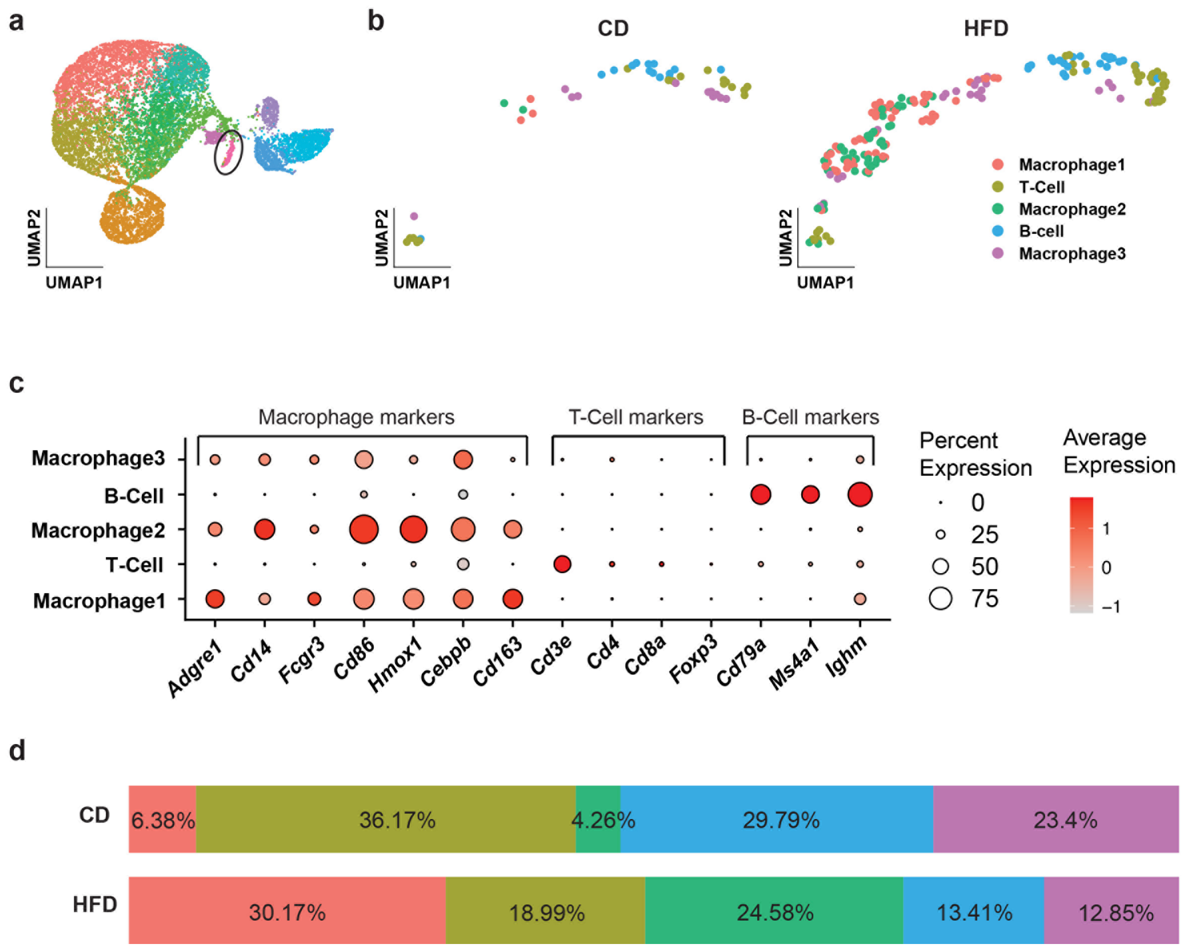


**Supplementary Figure 4. ScRNA-seq characterization of immune subpopulations in obesity. (a)** UMAP of total cells indicating selection of immune cluster. **(b)** UMAP of immune cells reclustered and split by diet condition, showing the presence of 3 macrophage, 1 T-cell, and 1 B-cell subpopulations. **(c)** Dot plot of gene expression indicating transcriptomic profile of immune subpopulations. **(d)** Bar plot comparison showing differences in distributions of immune subpopulations between HFD and CD conditions.
